# Deep Multimodal Representations and Classification of First-Episode Psychosis via Live Face Processing

**DOI:** 10.1101/2024.11.07.622469

**Authors:** Rahul Singh, Yanlei Zhang, Dhananjay Bhaskar, Vinod Srihari, Cenk Tek, Xian Zhang, J. Adam Noah, Smita Krishnaswamy, Joy Hirsch

## Abstract

Schizophrenia is a severe psychiatric disorder associated with a wide range of cognitive and neurophysiological dysfunctions and long-term social difficulties. In this paper, we test the hypothesis that integration of multiple simultaneous acquisitions of neuroimaging, behavioral, and clinical information will be better for prediction of early psychosis than unimodal recordings. We propose a novel framework to investigate the neural underpinnings of the early psychosis symptoms (that can develop into Schizophrenia with age) using multimodal acquisitions of neural and behavioral recordings including functional near-infrared spectroscopy (fNIRS) and electroencephalography (EEG), and facial features. Our data acquisition paradigm is based on live face-to-face interaction in order to study the neural correlates of social cognition in first-episode psychosis (FEP). We propose a novel deep representation learning framework, Neural-PRISM, for learning joint multimodal compressed representations combining neural as well as behavioral recordings. These learned representations are subsequently used to describe, classify, and predict the severity of early psychosis in patients, as measured by the Positive and Negative Syndrome Scale (PANSS) and Global Assessment of Functioning (GAF) scores. We found that incorporating joint multimodal representations from fNIRS and EEG along with behavioral recordings enhances classification between typical controls and FEP individuals. Additionally, our results suggest that geometric and topological features such as curvatures and path signatures of the embedded trajectories of brain activity enable detection of discriminatory neural characteristics in early psychosis.

## 1 Introduction

Schizophrenia is a complex mental disorder affecting millions of people worldwide. Individuals suffering from this condition face significant cognitive and social impairments. Current diagnostic methods, often based on static or single-subject studies, fail to capture the dynamic nature of social cognition, especially in interpreting facial expressions. This presents a challenge in early detection of the condition and subsequent early interventions that could improve the quality of life. Moreover, most existing methods focus on analyzing different neuroimaging and behavioral modalities separately, missing the intricate interactions between neural activities and their relationships to behavior. To address this, we propose a novel approach that combines live social interactions with multimodal neuroimaging (fNIRS, EEG) and facial expression analysis. Our method captures dynamic neural correlates of live face-to-face interactions in first-episode psychosis (FEP) patients, using a deep recurrent geometric autoencoder framework, that we call Neural-PRISM, to learn joint representations from these modalities, offering new insights and early predictive capabilities for clinical outcomes.

According to the Global Burden of Disease 2019 Study (Vos et al., 2020; Solmi et al., 2023), schizophrenia affects 23.6 million individuals worldwide. It is marked by positive symptoms such as delusions, hallucinations, and disorganized thinking, as well as negative symptoms including reduced speech, social withdrawal, and diminished emotional expression. The wide spectrum of cognitive and neurophysiological dysfunctions associated with Schizophrenia impose a profound impact on quality of life and social functioning. Moreover, the estimated economic burden of schizophrenia in the USA doubled from 2013 to 2019, reaching $343.2 billion in 2019 (Kadakia et al., 2022). This underscores the importance of developing effective early diagnosis strategies and treatment options to better manage this challenging disorder. However, studying schizophrenia using only unimodal neuroimaging or behavioral data is challenging because each offers a limited perspective, making it difficult to fully understand and address the cognitive and social deficits associated with the disorder. EEG offers high temporal but low spatial resolution, whereas, fNIRS provides better spatial but lower temporal resolution. Similarly, relying solely on behavioral data, like facial expression analysis, does not reveal the underlying neural mechanisms contributing to the observed impairments in schizophrenia. Some studies based on unimodal neuroimaging recordings include resting state functional magnetic resonance imaging (fMRI) (Cai et al., 2020; Li et al., 2020; Yassin et al., 2020; Lee et al., 2022) and resting state scalp electroencephalography (EEG) (Sun et al., 2021; Miras et al., 2023). Although schizophrenia is often associated with disordered social interactions, much of the current understanding of its underlying neurophysiology comes from studies of single brains without social interaction. To address this issue we focus on dynamic behavior during social interactions.

Recently, an emerging focus on live social interactions between pairs of individuals, rather than single subjects, has improved the understanding of dynamic face processing as a proxy for real-life social interactions (Noah et al., 2020; Hirsch, Zhang, Noah, Dravida, et al., 2022; Hirsch, Zhang, Noah, and Bhattacharya, 2023). These foundational findings provide a theoretical framework to study live face-to-face interactions in autism spectrum disorder (ASD) (Zhang et al., 2024), where social difficulties are a primary symptom. This research prompts new questions about atypical dynamic and interactive face processing as an indicator of underlying neurophysiology for social function and/or social disability in schizophrenia. We hypothesize that the neural systems of FEP patients as compared to TD individuals reflect characteristic atypical social functioning and suggest that they could serve as early indicators of risk, predictors of disease progression, and potential targets for interventions such as neuromodulation. Thus, here we apply this novel method of neural and behavioral recordings during live social interactions to isolate fundamental neural correlates characteristic of atypical social cognition in schizophrenia.

Functional magnetic resonance imaging (fMRI) provides high spatial but limited temporal resolution (approximately 2 seconds). However, fMRI is limited to single subject tasks, other constraining conditions, and a high magnetic field that limits simultaneous measurement of related behaviors. Functional near infrared spectroscopy, fNIRS, like fMRI also measures the hemodynamic response function (HRF) but at much higher temporal resolution. A limitation of fNIRS, relative to fMRI, is the shallow signal penetration that is restricted to superficial cortex. However, superficial cortical activity is assumed to reflect subcortical activity from deeper structures, and the fNIRS technology adds the key dimension of live behaviors within live social interactions. Thus, this limitation of responses to superficial cortex and relatively low spatial resolution is balanced with advantages of two-person social neuroscience behaviors that extend conventional single-subject neuroscience to dyadic functions and live reciprocal social interactions that cannot be observed using conventional neuroimaging methods. Here we apply live two-person interactive paradigm with *simultaneous* EEG and fNIRS recordings to investigate social cognitive mechanisms by live (ecologically valid) facial expressions (Wild et al., 2003) in both typically developing (TD) and FEP participants. These investigations are not possible with fMRI because live face-to-face imaging is difficult and the high magnetic field prevents incorporating other imaging modalities simultaneously.

To gain insights from this multimodal data, in this paper we propose a novel multimodal representation learning framework called neural-PRISM (Path Representations for early Identification of Schizophrenia via Multimodal translation) for extracting signatures of brain activity in FEP. The proposed neural-PRISM is a recurrent geometric autoencoder framework that learns compressed and informative latent representations of multiple modalities including fNIRS, EEG, and behavior in form of facial action units(AUs) (Ekman and Friesen, 1978). These representations reveal a highly structured and temporally organized trajectory in 3-D, with high-curvature segments corresponding to transitions in brain activity between live interactions and rest. Both encoder and decoder networks consist of multiple long short-term memory (LSTM) (Hochreiter and Schmidhuber, 1997; Goodfellow, Bengio, and Courville, 2016) layers to learn latent representations from neuroimaging (EEG and fNIRS) as well as behavioral (faceAU) modalities. These latent trajectories are utilized for distinguishing between FEP patients and typically developed (TD) individuals, as well as for forecasting clinical scores, such as Positive and Negative Syndrome Scale (PANSS) and Global Assessment of Functioning (GAF) scores (Jones et al., 1995; Srihari et al., 2015) that indicate the severity of psychosis. The learned representations are utilized via nonlinear dimensionality reduction method, t-PHATE (Busch et al., 2023), to visualize the neural activity in a three dimensional Euclidean space. We call these time lapse trajectories as neural motifs, which are further utilized for computing geometrical (curvatures) and topological (path signatures) features and discriminate between TD and FEP individuals.

To summarize, the contributions of this paper are as follows: (i) a novel live interactive paradigm with simultaneous fNIRS, EEG, and facial expression recordings to study the relationship between the neural correlates of FEP patients stimulated by social interaction and (ii) a novel recurrent geometric autoencoder framework called neural-PRISM for learning joint representations of multiple modalities. (iii) Empirical results demonstrating effective representation learning via visualization as well as classification result showing early FEP prediction.

## 2 Methods

### 2.1 Dataset and Experimental Setup

The proposed method employs dyads that include one individual who serves as the live expressive face stimulus and the other partner categorized as either typically developed (TD) or first episode psychosis (FEP) patient. Dyads faced each from across a table at a distance of approximately 140 cm and table-mounted eye-tracking systems were positioned to measure continuous eye movements of the subject. Functional NIRS and EEG data were also synchronized and continuously acquired hemodynamic and electrocortical responses of the subject during the experiment. The dyads were separated by a “smart glass” in the center of the table that controlled face gaze times (the glass was transparent during gaze periods) and “rest times” (the glass was opaque during rest) (Hirsch, Zhang, Noah, and Bhattacharya, 2023). The face gaze times were controlled according to the time series illustrated in Figure 1.

**Figure 1.**
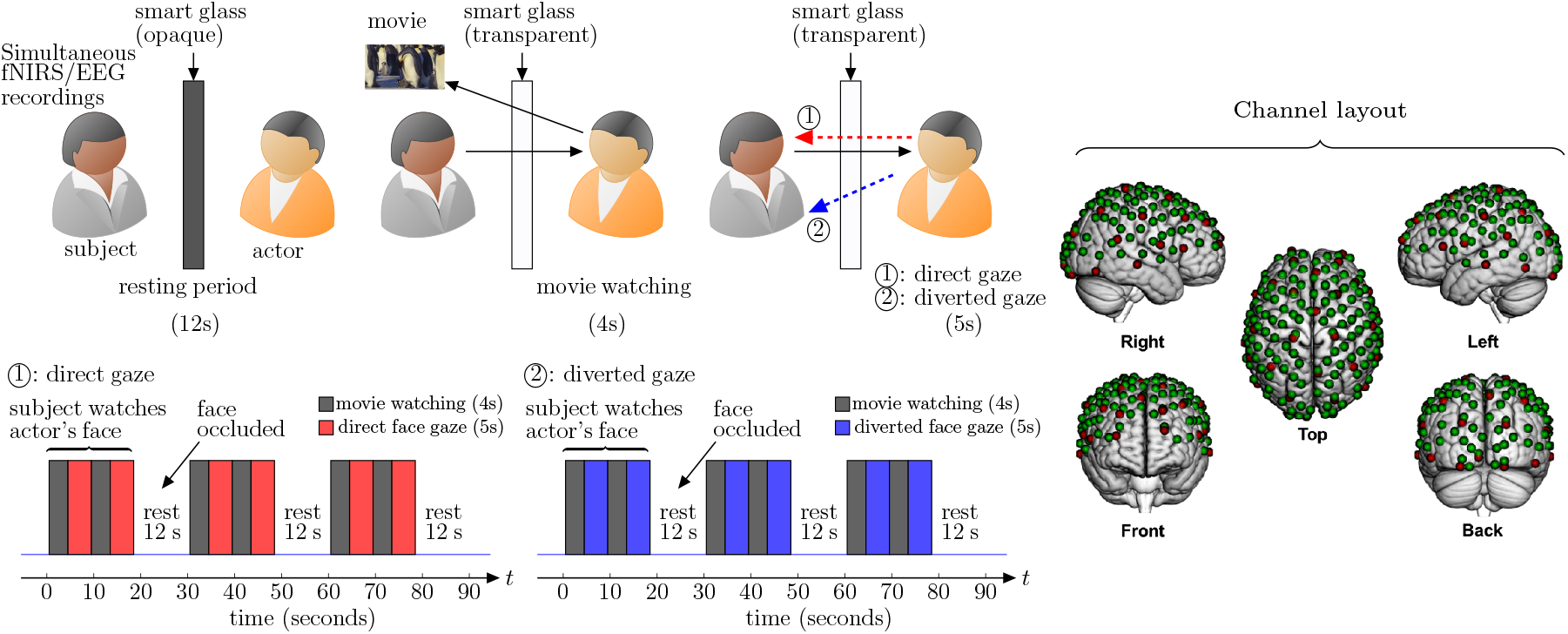
Experiment Setup: the subject’s brain is being scanned with simultaneous fNIRS, EEG, and facial expression recordings. The actor watches a (positive/negative) movie for 4 seconds followed by looking at the subject (eye contact/ no eye contact) for 5 seconds. The same process of 9 seconds is repeated again before the smart glass is made opaque for 12 seconds (rest period). This sequence of 30 seconds activity is repeated three times in a single run. Channel layout for simultaneous EEG and fNIRS recordings: red dots represent the 32 EEG electrodes and green dots represent the 134 fNIRS channels.

#### 2.1.1 Participants

Our study involved 14 FEP patients (2 females, 12 males; mean age: 24.2 ± 4.1 years) and 19 typical controls (8 females, 8 males and 3 identified as another gender; mean age: 25.1 ± 9.0 years). FEP patients were recruited from Connecticut Mental Health Center and Yale New Haven Hospital and the typically developing (TD) participants were recruited from the local community. All participants provided written informed consent in accordance with guidelines approved by the Yale University Human Investigation Committee (HIC # 1501015178).

#### 2.1.2 Paradigm

The dyads were seated 140 cm across a table from each other. A “Smart Glass” (glass that is capable of alternating its appearance between opaque and transparent upon application of an appropriate voltage) panel was positioned in the middle of the table 70 cm away from each participant. In both conditions of direct and diverted face gaze, the subject was instructed to gaze at the eyes of their partner who watches emotionally valanced movie clips followed by direct or diverted gaze towards the subjects face (Figure 1). In the direct face gaze condition, dyads had a direct face-to-face view of each other. On the other hand, in the diverted face gaze condition the stimulus look at the subject’s shoulder.

The actor watches a 4 second movie (joyful or sad) and then looks at the partner’s (subject’s) eyes or his shoulders (diverted face gaze) for 5 seconds. These sequence of tasks were repeated twice for each pair. Then there is a 12 second rest period, when the smart glass is made opaque. The same process (30 seconds) is repeated three times for each condition. The subjects were instructed to watch the actor’s (stimulus) face all the time. The actor was instructed to watch short movies followed by direct or diverted gaze towards the subject.

##### Movie Library

Emotionally evocative videos (movies) that are intended to elicit natural facial expressions were collected from publicly accessible sources and trimmed into 3-5 second clips. Video stimuli are pretested and rated for emotive properties along with 283 Amazon Mechanical Turk participants who rated 134 videos. The criteria for inclusion were that the videos be about 3-5 seconds in duration and have emotive inducing properties in accordance with three categories that we refer to as: adorables, creepies, and neutral landscapes. This is to avoid any presumption of emotional labels. This library of video clips has been employed previously to elicit dynamic and spontaneous facial expressions within a similar live-interaction paradigm (Hirsch, Zhang, Noah, and Bhattacharya, 2023). No video is repeated in any session.

#### 2.1.3 Functional Near-Infrared Spectroscopy Signal Acquisition

A Shimadzu LABNIRS system (Shimadzu Corp., Kyoto, Japan) was used to collect fNIRS data at a sampling rate of 123 ms (8.13 Hz). Forty emitters and forty detectors (80 optodes total) were placed in the cap in a 134-channel layout covering frontal, parietal, temporal, and occipital lobes (see channel layout in Figure 1) (Dravida et al., 2019). Each emitter transmitted three wavelengths of light, 780, 805, and 830 nm, and each detector measured the amount of light that was not absorbed. The amount of light absorbed by the blood was converted to concentrations of OxyHb and deOxyHb using the Beer-Lambert equation. Custom-made caps with interspersed optode and electrode holders were used to acquire concurrent fNIRS and EEG signals (Shimadzu Corp., Kyoto, Japan). The distance between optodes was 2.75 cm or 3 cm, respectively, for participants with head circumferences less than 56.5 cm or greater than 56.5 cm. Caps were placed such that the most anterior midline optode holder was almost 2.0 cm above nasion, and the most posterior and inferior midline optode holder was on or below inion. A lighted fibre-optic probe (Daiso, Hiroshima, Japan) was used to remove all hair from the optode holder before optode placement.

#### 2.1.4 Electroencephalograph Signal Acquisition

A g.USBamp (g.tec medical engineering GmbH, Austria) system with 2 bio-amplifiers and 32 electrodes was used to collect EEG data at a sampling rate of 256 Hz. Electrodes were arranged in a layout similar to the 10-10 system; however, exact positioning was limited by the location of the electrode holders, which were held rigid between the optode holders. Electrodes were placed as closely as possible to the following positions: Fp1, Fp2, AF3, AF4, F7, F3, Fz, F4, F8, PC5, PC1, PC2, PC6, T7, C3, Cz, C4, T8, CP5, CP1, CP2, CP6, P7, P3, Pz, P4, P8, PO3, PO4, O1, Oz, and O2. Conductive gel was applied to each electrode to reduce resistance by ensuring contact between the electrodes and the scalp. As gel was applied, data were visualized using a bandpass filter to allow frequencies between 1 and 60 Hz. The ground electrode was placed on the forehead between AF3 and AF4, and an ear clip was used for reference.

#### 2.1.5 Facial Features Acquisition

The behavioral data for the subjects was simultaneously acquired in form of facial action units (AUs) using OpenFace (Baltrušaitis, Robinson, and Morency, 2016) and Logitech C920 face cameras. OpenFace is one of several available platforms that provide algorithmically derived tracking of facial motion in both binary and continuous format. The automatic detection of facial AUs using these tools has become a foundational method in facial expression analysis, where facial movements are characterized as dynamic patterns reflecting the anatomy of facial muscles. While a direct link between specific emotions and activation patterns has been proposed (Ekman, 1993), this approach focuses on breaking down facial expressions into discrete muscular components and their dynamics, without associating them with emotional labels. The facial AU data from OpenFace included 17 distinct classifications of anatomical configurations.

### 2.2 Representation Learning and Classification

With the experimental setup for data collection discussed in the previous section, we propose a novel deep recurrent geometric autoencoder framework, neural-PRISM (Path Representations for early Identification of Schizophrenia via Multimodal translation), for learning unimodal and multimodal (joint) representations. We call these learned representations as “neural motifs” which represent the underlying signatures of timelapse neuroimaging and/or behavioral data in a compressed low-dimensional space.

#### 2.2.1 Latent Trajectories from Multimodal Translation via Recurrent Autoencoders

Autoencoders are powerful machine learning models trained in a self-supervised fashion to reconstruct inputs by learning their abstract representations in the latent space. Besides learning representations for a single modality, the encoder decoder framework can also be utilized for learning joint representations of two neuroimaging modalities with input to the encoder being data from one modality and the decoder output being the other modality (Figure 2a). We utilize deep recurrent neural networks (RNNs) (Hermans and Schrauwen, 2013) for encoder as well as decoder to capture the time dependency of the neural recordings and facial action units. Both the encoder and decoder networks consist of RNN layers: the encoder network encodes the multidimensional input neural recordings (EEG or fNIRS) **x** into latent representations

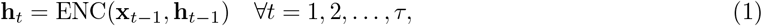

where *τ* is the length of the input sequence. The dimensions of the input at each time-point is **x**_*t*_ *∈* ℝ^134^ for fNIRS or facial action units and **x**_*t*_ *∈* ℝ^32^ for EEG. The latent embeddings **h**_*t*_ represent the compressed time encoded information in the input. The final latent embedding **h**_*τ*_ encapsulates the temporal patterns present in the input and serves as the initialization for the decoder. The decoder network takes the latent embeddings **h**_*τ*_ and generates the reconstructed data, with its outputs computed at each time step *t*

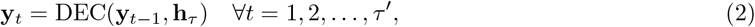

where *τ*′ is the sequence length of the Decoder output modality. In our settings, the EEG data has 7680 samples corresponding to the 30 second block while fNIRS and face AU constitute 244 samples. Note that we consider HbDiff signal (Kirilina et al., 2012), difference between OxyHb and deOxyHb signals, as fNIRS recordings.

**Figure 2.**
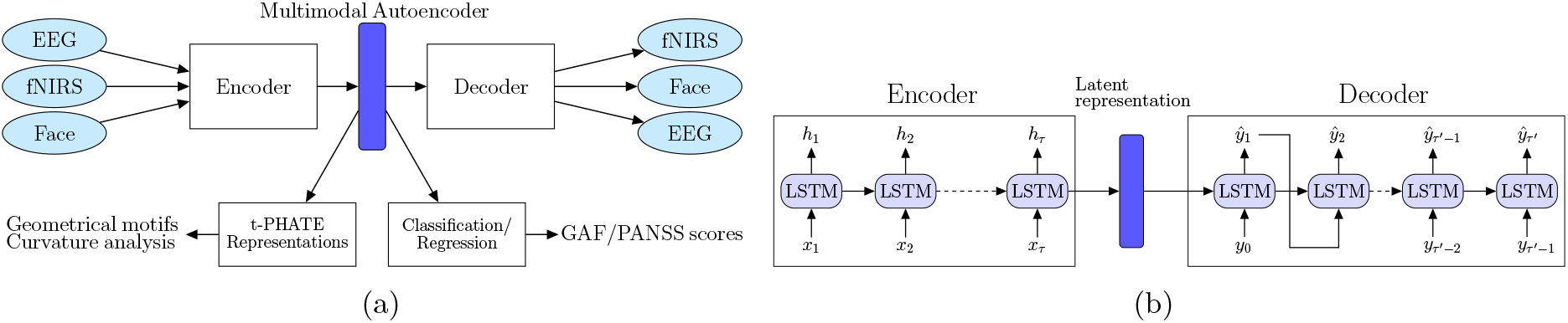
(a) Schematic of our neural-PRISM recurrent geometric autoencoder framework. (b) Architecture of encoder and decoder networks.

A Long-Short-Term-Memory (LSTM) RNN, as shown in Figure 2b, was chosen over the vanilla RNN because the latter experiences the vanishing-gradient problem during model training, which inhibited it from effectively leveraging context between elements by maintaining its internal state. For the decoder network, teacher forcing method (Williams and Zipser, 1989; Lamb et al., 2016) was employed, in which the groundtruth samples **y**_*t*_ are fed back into the model to be conditioned on for the prediction of later outputs. These fed back samples force the RNN to stay close to the ground-truth sequence.

The final latent embeddings **h**_*τ*_ are fed to a multilayer perceptron (Goodfellow, Bengio, and Courville, 2016) layer in order to classify FEP vs TD individuals. The learned trajectories (**h**_1_, **h**_2_, …, **h**_*τ*_) in the latent space are further analyzed topologically and geometrically, as described in the following section. Our recurrent geometric autoencoder framework also offers a foundational approach for translating between different modalities. While other studies, such as (Sirpal et al., 2022), have focused solely on modality translation using resting-state EEG and fNIRS data, our primary goal here is not to advance translation techniques. Instead, the translation between neuroimaging modalities (EEG and fNIRS) and behavioral modalities (FaceAU) is an additional outcome of our framework.

#### 2.2.2 Topological and Geometrical Summarization of Latent Trajectories

We summarize the high-dimensional latent trajectories obtained from the recurrent autoencoder using path signatures, subsequently leveraging these path signatures for classification. Path signatures (Chevyrev and Kormilitzin, 2016), as effective descriptors of ordered data, capture essential characteristics of trajectories and have been successfully applied in various domains of neuroscience. For instance, path signatures have been employed to predict Alzheimer’s diagnosis by modeling disease progression trajectories (Moore et al., 2019), to detect epileptic seizures by analyzing electroencephalogram (EEG) patterns (Tang et al., 2024), in early autism diagnosis through behavioral pattern recognition (Yin et al., 2024), and in seizure forecasting (Haderlein et al., 2023).

Next, we reduce the dimensionality of the latent representations using the manifold learning technique tPHATE (Busch et al., 2023). tPHATE preserves local and global structures in the data, while simultaneously enabling us to visualize it in 3-D. By embedding the high-dimensional latent trajectories into a lower-dimensional space, we can compute and analyze the geometric features of the resulting low-dimensional trajectories, such as curvature. Here we employ curvature as a feature for classification, as it encapsulates information about changes in trajectory direction. Curvature analysis of dynamic trajectories has been widely used in scientific machine learning, including shape analysis in computer vision (Coeurjolly, Miguet, and Tougne, 2001), understanding particle movement in physics (Thiel, 2024), and analyzing motor control and movement dynamics in neuroscience (Tschechne and Neumann, 2014; Rocchi et al., 2007).

Overall, our approach of using geometrical and topological summaries of latent trajectories (Figure 3), described below, enables a nuanced classification framework that leverages both temporal ordering and geometric properties of brain activity.

**Figure 3.**
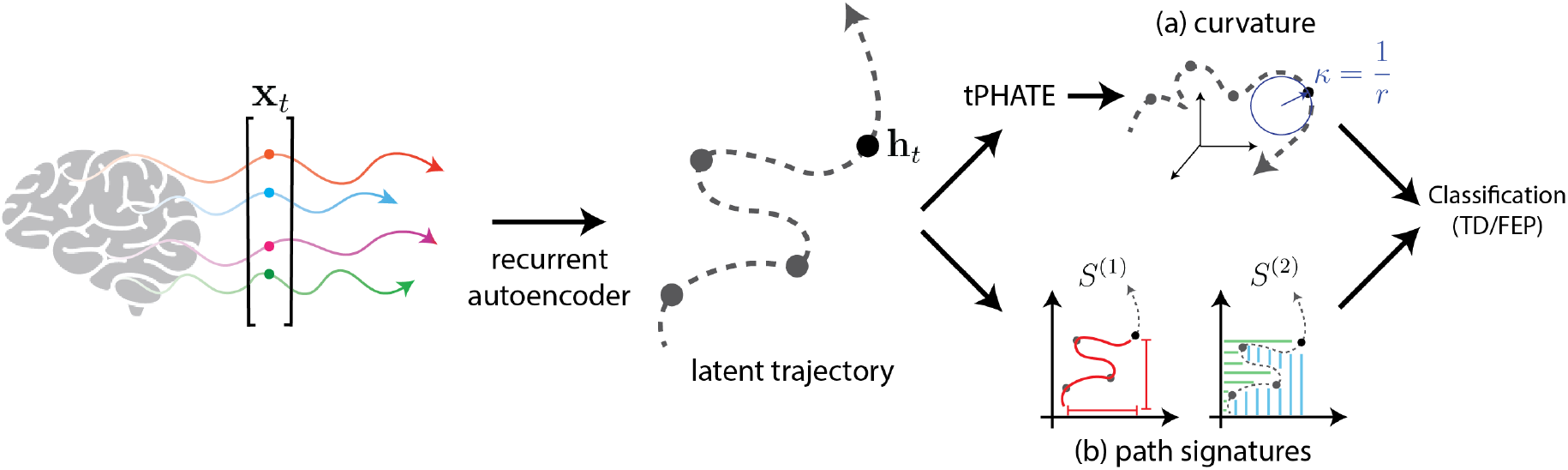
Classification of subjects based on curvature and path signatures of the latent trajectories obtained from the recurrent autoencoder. (a) Curvature computation is performed using circle fitting in 3D tPHATE coordinates. (b) Path signatures are directly computed from the latent trajectory.

##### Path Signatures

Given the latent trajectory **h**(*t*) = (**h**_1_, **h**_2_, …, **h**_*τ*_), each dimension *h*_*i*_(*t*) = (*h*_*i*1_, *h*_*i*2_, …, *h*_*iτ*_), is first rescaled to unit variance, reducing scale discrepancies among features. This is accomplished by standardizing each component of the path **h**(*t*) as follows:

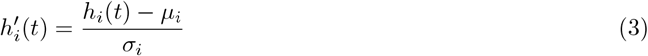

where *μ*_*i*_ and *σ*_*i*_ are the mean and standard deviation, respectively, of the *i*-th component across all time points. This normalization step ensures that each dimension contributes equally to the signature computation, minimizing bias toward features with larger scales.

To further address variability in the duration and sampling intervals across different modalities, we apply a time rescaling that standardizes the time interval of analysis. Specifically, we transform the time interval of interest [*a, b*] to the standard interval [0, 1]:

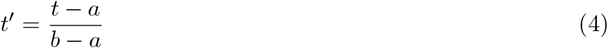

Following these preprocessing steps, we compute the *k*-th level path signatures, *S*^*k*^(**h**′(*t*)) for *k* = 1, …, *N* (see Appendix A for details). Each computed signature is then normalized:

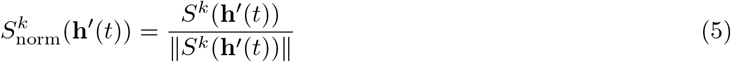

The normalized path signatures are subsequently fed into a four-layer multilayer perceptron (MLP) for classification.

##### PHATE and t-PHATE

Traditional dimensionality-reduction techniques such as PCA, t-SNE (Van der Maaten and Hinton, 2008) and UMAP (McInnes, Healy, and Melville, 2018) are suboptimal: they are sensitive to noise, scramble global structures, fail to capture fine-grained local details, and often lack scalability for large datasets (Moon et al., 2019). To overcome these challenges, PHATE (potential of heat diffusion for affinity-based transition embedding) (Moon et al., 2019) provides a scalable dimensionality-reduction method that gives accurate, denoised visualizations of both local and global structures without imposing strong structural assumptions.

By incorporating time-varying features, t-PHATE (Busch et al., 2023) extends the PHATE algorithm to model the temporal properties of input signals, capturing both temporal autocorrelation and stimulus-specific dynamics. When applied to fMRI data from cognitive tasks, it denoises the data and enhances access to brain-state trajectories compared to voxel data and other embeddings like PCA, UMAP, t-SNE and PHATE. Through the integration of temporal relationships between LSTM cells at different time points, t-PHATE generates a low-dimensional (3-D) embeddings that capture both the spatial organization of LSTM states and their temporal progression.

##### Geometrical Feature Extraction

One observation from the t-PHATE embeddings is that the rate of directional change over time in each trajectory correlates with the intensity of attention shifts during task-switch periods (see Figure 5a). This insight motivates the further use of three-dimensional t-PHATE embeddings for feature extraction in the form of curvature measures. More precisely, the curvature at a specific point reflects the rate of change of the curve at that point, or in mathematical terms, it represents the magnitude of the second derivative of the curve at that point. A plane curve given by Cartesian parametric equations *x* = *x*(*t*) and *y* = *y*(*t*), the curvature kappa, sometimes also called the “first curvature”(Kreyszig, 1991), is defined by

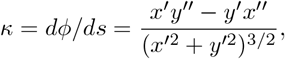

where *x*′ and *x*^*′′*^ denote first and second order derivatives, respectively. Here we consider 1-dimensional curves in 3-dimensional Euclidean space, specified parametrically by *x* = *r* cos *t* and *y* = *r* sin *t*, which is tangent to the curve at a given point. The curvature is then

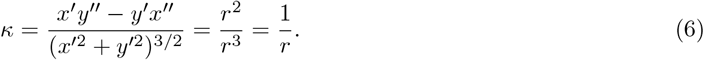

For curvature at point *p*, we fit a circle *S*^1^(*p, r*) centered at *p* with radius *r* in the plane spanned by principal components of the 3-dimensional t-PHATE trajectory. The inverse of radius 1*/r* gives the curvature at *p*. More precisely, at each point *p* of the curve *C*, we select a local neighborhood of points around *p*. The size of this neighborhood, a user-defined hyper-parameter (set here to 8% of the total curve length), determines the number of points sampled symmetrically around *p*. The neighborhood is then centered by subtracting the mean of these points from each point, ensuring that the analysis is performed relative to the center of mass. Next, Singular Value Decomposition (SVD) is applied to the centered neighborhood, yielding two vectors that span the local plane and a normal vector perpendicular to this plane. A circle is then fitted to the points in the local plane using a least-squares method. The curvature at *p* is subsequently calculated as the reciprocal of the radius (1*/r*) of the fitted circle, assuming that locally the trajectory approximates a circular arc. This procedure is repeated for all points along the trajectory, giving a curvature profile across the entire curve. The four curvature values at four task switching times are then selected to be fed into a three-layer MLP for classification. Detailed classification accuracies can be found in Table 1 in Appendix.

**Table 1:** Classification accuracies based on curvature of tPHATE embeddings and path signatures obtained from unimodal and multimodal latent trajectories generated by neural-PRISM with input from various EEG bands.

| Feature | EEG Band | Unimodal | Multimodal |  |
| --- | --- | --- | --- | --- |
|  |  |  | EEG + fNIRS | EEG + FaceAU |
| Curvature | alpha | 0.572 | 0.617 | 0.632 |
|  | delta | 0.619 | 0.672 | 0.635 |
|  | theta | **0.872** | 0.677 | 0.636 |
| Path Signature | alpha | 0.693 | 0.826 | 0.839 |
|  | delta | 0.654 | 0.893 | 0.823 |
|  | theta | 0.722 | **0.917** | **0.871** |

## 3 Results

We present our experimental results in two parts. First, we present the classification results from learned representations followed by prediction of GAF and PANSS scores. Next, we present the joint learned representations and show the distinction between TD and FEP individuals via geometrical data analysis techniques.

### 3.1 Classification

We divide the dataset into 30 seconds blocks such that each subject has 24 blocks of data: with positive/negative valence movies and direct/diverted gaze, corresponding to each condition, we have 6 blocks. In order to evaluate the performance of our method, we employ leave-one-subject-out cross validation scheme, the samples from one subject are used for testing, while samples from other subjects are used as the training set. It has to noted here that if we randomly choose certain blocks from all the data samples and split it into training and test sets, we achieve close to ideal 100% classification accuracy on test samples (with fNIRS recordings only) similar to the studies in previous works (Sun et al., 2021; Miras et al., 2023). This is because the training set has some cues or signatures of every subject and as a consequence the leaned model is able to generalize in this setting.

We train the encoder-decoder model with different EEG bands namely: delta [0.5–3 Hz], theta [4–7 Hz], and alpha [8–13 Hz] along with fNIRS and facial action units. The classification accuracy achieved using fNIRS data on withheld subject blocks is determined to be 85 %, outperforming traditional support vector machine (SVM) accuracy of 71% and stand-alone MLP accuracy of 67%. Incorporating multimodal joint representations improves the classification and fNIRS + EEG data yields best classification accuracy of 88% (Figure 4(a)).

**Figure 4.**
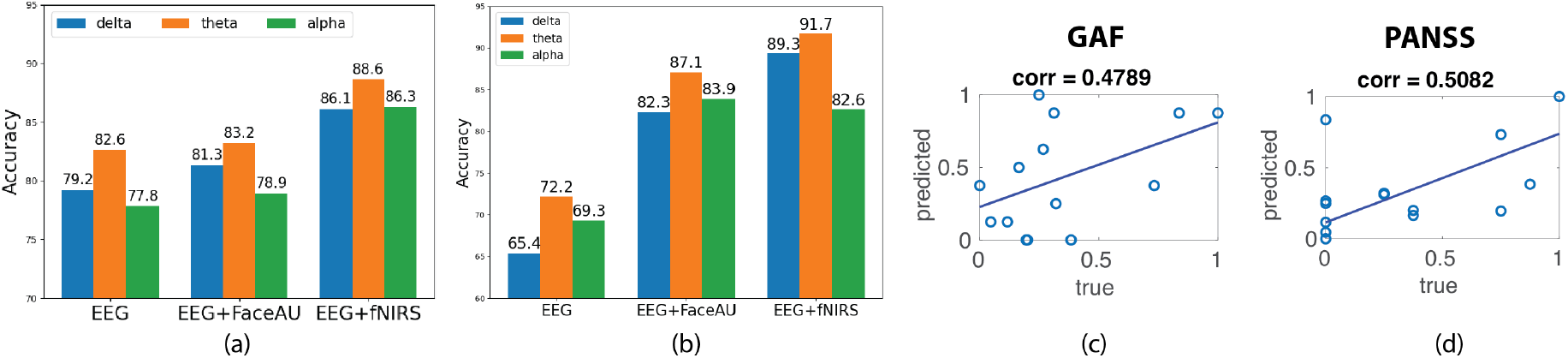
Diagnosis of FEP patients using latent trajectories obtained from neural-PRISM and prediction of disease severity scores. (a) Classification of TD and FEP subjects using a 2-layer multi-layer perceptron (MLP) trained on latent trajectories derived from unimodal (EEG) and multimodal (EEG+FaceAU and EEG+fNIRS) data. (b) Classification of TD and FEP subjects using a 4-layer MLP trained on path signatures of latent trajectories derived from unimodal and multimodal data. (c) Block-averaged probability scores obtained from the classifier in (a) are correlated with GAF scores. (d) Block-averaged probability scores obtained from the classifier in (a) are correlated with PANSS positive symptom score. Note that the PANSS and GAF scores are normalized to [0,1] for clarity.

#### 3.1.1 Predicting GAF and PANSS Scores

The Global Assessment of Functioning (GAF) (Aas, 2010; Srihari et al., 2015) covers the range from positive mental health to severe psychopathology, is an overall (global) measure of how patients are doing in their day-to-day life. GAF measures the degree of mental illness by rating psycho-logical, social and occupational functioning (Söderberg, Tungström, and Armelius, 2005). The Positive and Negative Syndrome Scale (PANSS) (Kay, Fiszbein, and Opler, 1987; Leucht et al., 2005) was developed in order to provide a well-defined instrument to specifically assess both positive and negative symptoms of schizophrenia as well as general psychopathology.

The classification probability scores during testing of classification model were utilized to predict the PANSS and GAF scores. The probability scores corresponding to the 24 blocks of data for each FEP patient were averaged to get the predicted score. Note that the ground truth scores were not used during training of our classification model. The correlation coefficient between predicted scores and true GAF role scores is computed at 0.4789 while correlation between positive symptom PANSS score was 0.5082 (Figure 4 (b) and (c)). However, the predicted scores did not have good correlation with the negative PANSS scores.

### 3.2 Learned Representations

The learned latent representations of the unimodal and multimodal autoencoders are used to compute the time lapse t-PHATE trajectories. We subsequently analyze these trajectories and compute curvatures at different task switching times. We observe that the curvatures for FEP patients are larger than those for TD individuals, indicating the attentional dysregulation and sensitivity to the presence of emotional distractors in FEP patients (Nestor and O’Donnell, 1998; Grave et al., 2023). Additionally, visualizing the learned embeddings in 3-D space using t-PHATE enables the identification of task switching times from movie watching to direct/diverted gaze and vice-versa (Figure 5(a) shows example trajectories). The curvatures at various switching times were analyzed for both FEP and TD individuals. By computing curvatures using only EEG unimodal representations, the greatest distinction between TD and FEP individuals was observed in the theta band (Figure 5(b)). Moreover, by integrating different modalities - fNIRS and FaceAU - with EEG, clear discrimination emerges in both the alpha and delta bands (Figure 5(c) and (d)).

**Figure 5.**
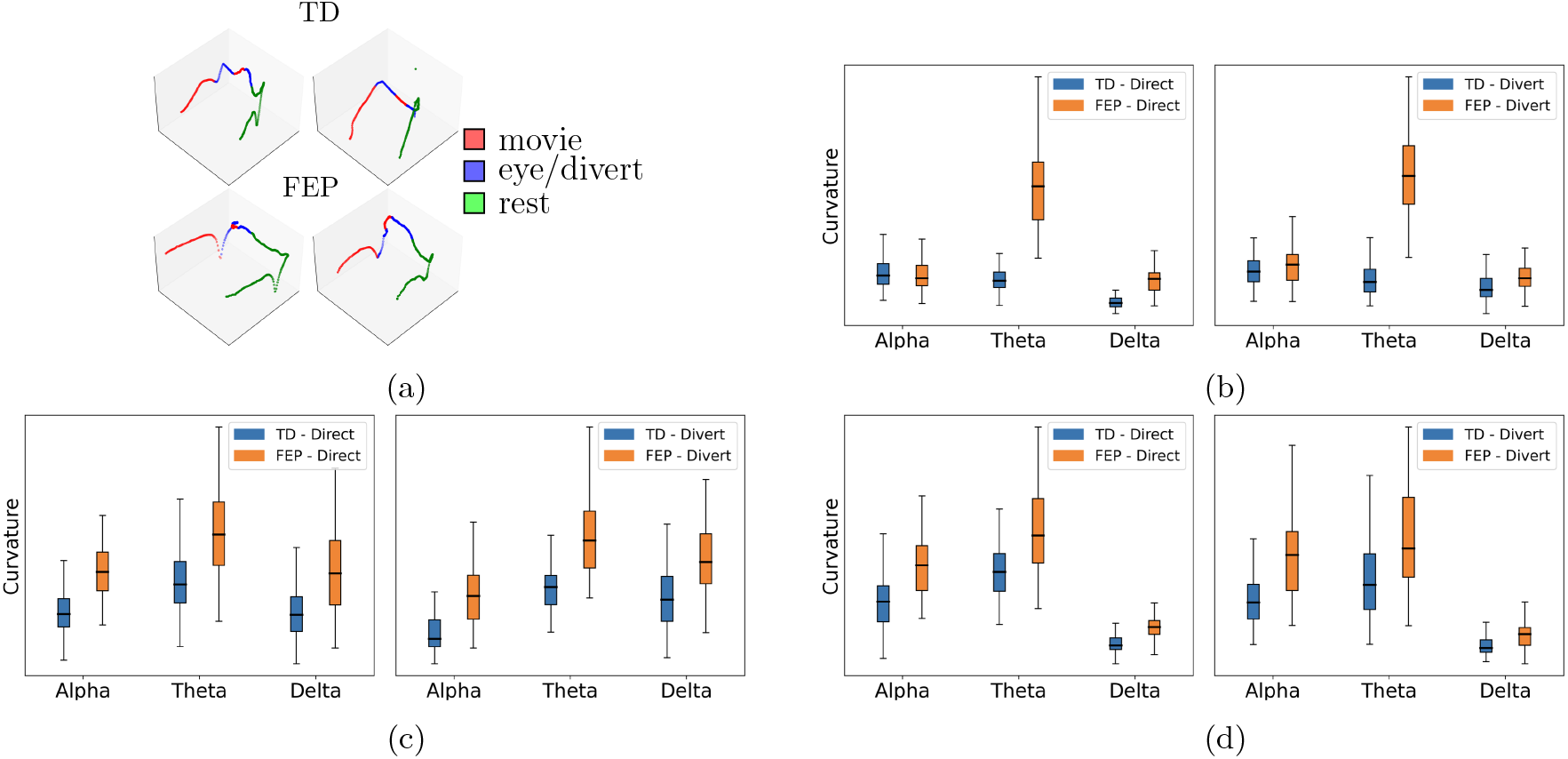
(a) Example t-PHATE visualizations (geometrical motifs) of learned representations in 3-dimensional Euclidean space: first row is for TD and second row corresponds to FEP patients. These examples are obtained from joint EEG and fNIRS representations. Curvatures of geometrical motifs: (b) unimodal EEG, (c) Multimodal EEG + FaceAU, and (d) Multimodal EEG + fNIRS.

## Discussion

In this study, we introduced a deep recurrent geometric autoencoder framework for multimodal representation learning and classification of first episode psychosis individuals. Our study is based on a live face-to-face interaction paradigm to investigate the neural correlates of social cognition in early psychosis. We hypothesized that incorporating multiple neuroimaging modalities (fNIRS and EEG) along with behavioral recordings (facial tracking) can predict early psychosis symptoms better than unimodal recordings alone.

Our proposed neural-PRISM framework consists of LSTM based encoder and decoder networks together with geometric and topological characterizations of the learned trajectories. The encoder network is trained to output latent trajectories over time and the decoder network is trained to output reconstructed modality conditioned on the encoder output. By training the networks to minimize the difference between the predicted and recorded (ground truth) output modalities, the autoencoder learns compressed joint embeddings of multimodal neural trajectories in the latent space (encoder output). Although the classification between FEP and TD individuals is based on the embedding at the final time points, the entire learned trajectories are utilized to capture geometrical features that facilitate characterization of early psychosis patients.

The classification and severity prediction results in Figure 4 as trajectory curvature analysis on our embeddings results in Figure 5 support our hypothesis, where the multimodal representations provide better discrimination for FEP patients. The curvatures of trajectories associated with the task-switching paradigm may indicate rapid transitions between events, which could be reflected in the EEG data which has higher temporal resolution. Moreover, higher curvatures of FEP patients validate the attentional dysregulation and sensitivity to the presence of emotional distractors in FEP patients (Nestor and O’Donnell, 1998; Grave et al., 2023), underlying the validity of our approach.

Our results confirm the potential of our framework for facilitating classification and detection of early psychosis. We achieve higher classification accuracy from multimodal (EEG and fNIRS/FaceAU) joint representation learning than achieved solely from fNIRS data validating the usefulness of multimodal data processing. Moreover, our paradigm along with multivariate data analysis show correlations with early positive symptoms and this may aid clinicians at targeting for intervention.

Although the number of participants in the study is small, the current set of data provides foundational results with multivariate analysis techniques for potential future studies on larger populations as well as application of these tools to additional populations including chronic schizophrenia. Further generalization across subjects will require a larger sample size with a primary emphasis on understanding FEP through neural recordings stimulated by live face processing.

While fNIRS has been extensively used for neuroimaging in infants and children, its application in adult cognitive research has been limited, primarily due to sparse optode coverage and lower spatial resolution (around 3 cm) compared to fMRI. Nevertheless, its advantages, such as tolerance to movement and the absence of factors like a strong magnetic field, restrictive physical conditions, the requirement to lie supine, and loud noise, make it a preferable alternative for live interactive studies involving two individuals (Noah et al., 2020; Hirsch, Zhang, Noah, Dravida, et al., 2022; Hirsch, Zhang, Noah, and Bhattacharya, 2023). Although fNIRS technology cannot record brain activity from subcortical regions, many studies on social interaction have found the superficial cortex including right temporoparietal junction to play a major role in these behaviors (Carter and Huettel, 2013). Combining fNIRS recordings with EEG provides additional information that may represent neural processing at deeper and subcortical levels.

In conclusion, this study demonstrated the potential of multivariate techniques to capture discriminatory patterns in neural and behavioral recordings of early psychosis. Our findings provide a foundation for exploring the mechanisms underlying these conditions and their interconnections.

## Author Contributions

RS: Methodology, Visualization, Manuscript preparation. YZ: Methodology, Visualization, Manuscript preparation. DB: Methodology, Visualization, Manuscript preparation. VS: Clinical assessment & recruiting subjects, Manuscript preparation. CT: Clinical assessment & recruiting subjects, Manuscript preparation. XZ: Data acquisition & preprocessing, Manuscript preparation. AN: Data acquisition & preprocessing, Manuscript preparation. SK: Conceptualization - Computational methods, Funding acquisition, Manuscript preparation. JH: Conceptualization - Experimental paradigm & Data acquisition, Funding acquisition, Manuscript preparation.

## Acknowledgments

We thank Nina Levine, LMSW, MPH, and Deepa Purushothaman, MD, for their assistance in scanning FEP participants. We extend our sincere appreciation to Raymond Cappiello, PhD, for managing all the required documentation.

## Data Availability

The raw data supporting the conclusions of this article will be made available by the authors, without undue reservation.

## Code Availability

Our code will be made available on GitHub at: https://github.com/KrishnaswamyLab/neural-PRISM

## A Path Signatures

The path signature is a structured summary of a path in a multidimensional space, characterizing its properties by capturing all iterated integrals of the path components up to a specified truncation order. Following the formalism presented by Chevyrev and Kormilitzin (Chevyrev and Kormilitzin, 2016), given a piecewise differentiable path *X* : [*a, b*] → ℝ^*d*^, we define the signature of *X*, denoted *S*(*X*)_*a,b*_, is an infinite sequence where each term is derived from the iterated line integrals of *X*. Specifically, the *k*-th level signature, *S*^(*k*)^(*X*)_*a,b*_, is given by:

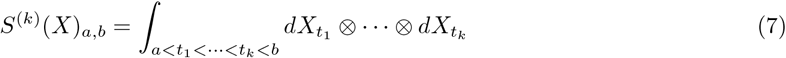

This sequence starts from *k* = 1 and continues indefinitely, capturing increasingly complex interactions among the path components over its domain. To compute this in practice, especially for paths represented by discrete data points 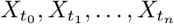 with *t*_0_ = *a* and *t*_*n*_ = *b*, the iterated integrals are approximated by summing over all ordered combinations of the sampled points:

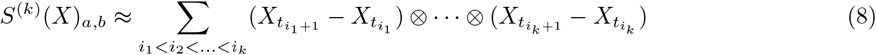

In order to manage computational costs, the signature is typically truncated at a finite level *N*, providing a compact yet informative representation:

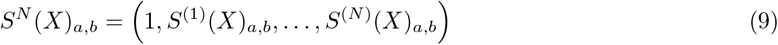

Each term of *S*^*N*^ (*X*)_*a,b*_ is a tensor product of differences between successive path points, efficiently encapsulating the path’s essential geometric and dynamic features up to the truncation level.

## B Implementation Details

We used PyTorch for our implementation. Three LSTM layers were used in both the encoder and decoder networks, with a latent dimension of 128. In the learning process, we utilized root mean square error (RMSE) as a loss function for training encoder and decoder networks, while cross entropy loss Mao, Mohri, and Zhong, 2023 was used for training classification models. Adam optimizer Kingma and Ba, 2015 is used along with *ℓ*_2_ regularization to prevent overfitting. The learning rate and weight decay (*ℓ*_2_ regularization) hyperparameters are tuned through grid search.

